# Inferring circadian rhythm disruptions in cancer using phase differences between clock genes

**DOI:** 10.1101/2025.01.18.633712

**Authors:** Ziang Zhang

## Abstract

Circadian rhythms exist across various levels of biological activities, from molecular processes to behaviors. Circadian clock genes are closely linked to the onset and progression of cancer; however, systematic analysis of their rhythmic expression in human tumors is still lacking due to difficulties in time-series sampling. In this study, we examined and improved the method for inferring phase differences through clock gene co-expression to investigate circadian rhythms in timestamp-free samples. We found that the co-expression levels of rhythmic genes are primarily determined by phase differences and are influenced by the strength and tissue specificity of rhythmic expression. Thus, we identified evolutionarily conserved rhythmic genes across multiple tissues to construct tissue-specific phase difference matrices. On this basis, we developed a method for inferring phase difference variations and extended it to the single-sample level as ssDistance to evaluate circadian rhythm disruptions in tumors. Results revealed that the significant alterations in phase differences between clock genes in tumors are cancer-type-specific and have complex effects on patient survival. This method provides an effective tool for studying circadian rhythms in large-scale public datasets lacking temporal information.

## Introduction

Circadian rhythms allow organisms to adjust to periodic environmental fluctuations resulting from Earth’s rotation^1^. At the molecular level, this endogenous rhythm is sustained by a network of interwoven positive and negative feedback loops^2,3^. In mammalian cells, the feedback loop’s activators primarily consist of *Arntl* (also referred to as *Bmal1*), *Clock*, and *Npas2*, which contain bHLH-PAS domains, while repressors include Per1, *Per2*, *Per3*, along with *Cry1* and *Cry2*^3,4^. Upon entering the nucleus, activators bind to cis-regulatory elements such as the E-box or D-box, initiating the transcription of repressors^3,4^. Subsequently, repressors inhibit activator function through protein-protein interactions or direct binding to cis-elements, until they are degraded via ubiquitination or other pathways^2–5^. Genes such as *Dbp*, *Nfil3*, *Nr1d1*, *Nr1d2*, *Rora*, and *Rorb* also participate in this network in a similar manner^4,6^. The rhythmic expression of these clock genes during the circadian cycle can be mathematically modeled using sine functions^2,5,7^.

Circadian rhythms have been found to be closely associated with cancer^8–12^. Genome-wide association studies have revealed a highly significant statistical association between genetic variations in clock genes and an increased risk of cancers such as breast, prostate, and lung cancer^9,13^. However, certain genetic variants of clock genes have been identified as having protective roles against cancer^9,13^. For instance, *CLOCK* polymorphisms rs11943456 and rs3749474 are significantly associated with an increased and decreased risk of breast cancer, respectively, while the *CRY1* variant rs1056560 is linked to a reduced risk of breast cancer in premenopausal women^9,14,15^. Pan-cancer transcriptomic analyses have also revealed widespread transcriptional dysregulation of clock genes in tumors^16–19^. Studies based on The Cancer Genome Atlas (TCGA) have shown that 90% of clock genes are differentially expressed in at least one cancer type, with certain genes exhibiting cancer-type-specific expression patterns^16^. For example, *PER2* is generally downregulated in most cancer types but upregulated in clear cell renal carcinoma^16^.

Despite these advances, the understanding of circadian rhythms in tumors of cancer patients remains limited, as it is impractical to perform multiple tumor tissue samplings within a 24-hour period, and most existing human cancer datasets lack circadian time information for sampling^13,20,21^. To address these limitations, several computational methods have been developed to analyze circadian rhythms in timestamp-free samples. One class of methods constructs scores based on the expression levels of clock genes to assess the activity of core clock gene expression, but these methods lack descriptions of circadian rhythm parameters^17,22–24^. Another class of methods infers the circadian time of sampling based on prior knowledge, which usually requires a large sample size^21,25–27^. A third class of methods uses the co-expression levels between clock genes as proxy variables for phase differences, assessing circadian rhythm disruptions in tumors by comparing clock gene co-expression patterns between normal and tumor tissues^20,28^. Studies employing the third type of method have revealed widespread alterations in clock gene co-expression across different cancer types^20^. Nevertheless, these methods still have aspects that can be further refined. Existing studies lack detailed examinations of the conversion between rhythmic gene co-expression levels and phase differences. Furthermore, there is a lack of rigorous selection and validation of genes used for matrix construction, with classical clock genes being manually chosen. Moreover, the reference matrices are not tissue-specific. Lastly, current analytical methods yield a single value for a group of samples, lacking resolution at the single-sample level.

In response to the limitations of the aforementioned methods based on clock gene co-expression, this study performed a more comprehensive analysis and refinement. First, using multi-tissue diurnal transcriptomic data from mice, we demonstrated that the co-expression levels between rhythmic genes are well accounted for by phase differences, and discovered that the strength and tissue specificity of rhythmic expression are key influencing factors in this explanatory power. Next, we identified evolutionarily conserved multi-tissue rhythmic genes in both mice and humans and derived tissue-specific phase difference (φ) matrices. Based on this, we developed a method for inferring phase difference variations (Δφ) and applied it to TCGA data, uncovering cancer-type-specific alterations in clock gene phase differences across multiple cancers. Finally, to extend this method to the single-sample level, we developed a circadian rhythm score based on phase differences, ssDistance. This score stratified cancer patients into subgroups with significantly different overall survival, revealing the dual role of clock gene phase differences in cancer risk.

## Results

### The co-expression of rhythmic genes is mainly governed by phase differences

The rhythmic expression of genes is generally characterized by sine functions, where the correlation between sine functions corresponds to the cosine of the phase difference^20,29–31^. Thus, the co-expression level (r) between rhythmic genes and the phase difference (φ) can be mutually converted (Figure 1A). This method is used to infer phase differences between clock genes in timestamp-free data^28^. However, a thorough evaluation of this application using real-world data is still lacking. To address this, we calculated the co-expression levels and phase differences between all rhythmic genes in each tissue using multi-tissue diurnal transcriptomic data from mice^32^. These were considered observed values, from which we derived corresponding predicted values, and we evaluated the consistency and absolute error between observed and predicted values (Figure 1B and 1C). Here, we define consistency as the Pearson correlation coefficient between a set of observed and predicted values, which should be distinguished from the correlation coefficient of co-expression.

**Figure 1.**
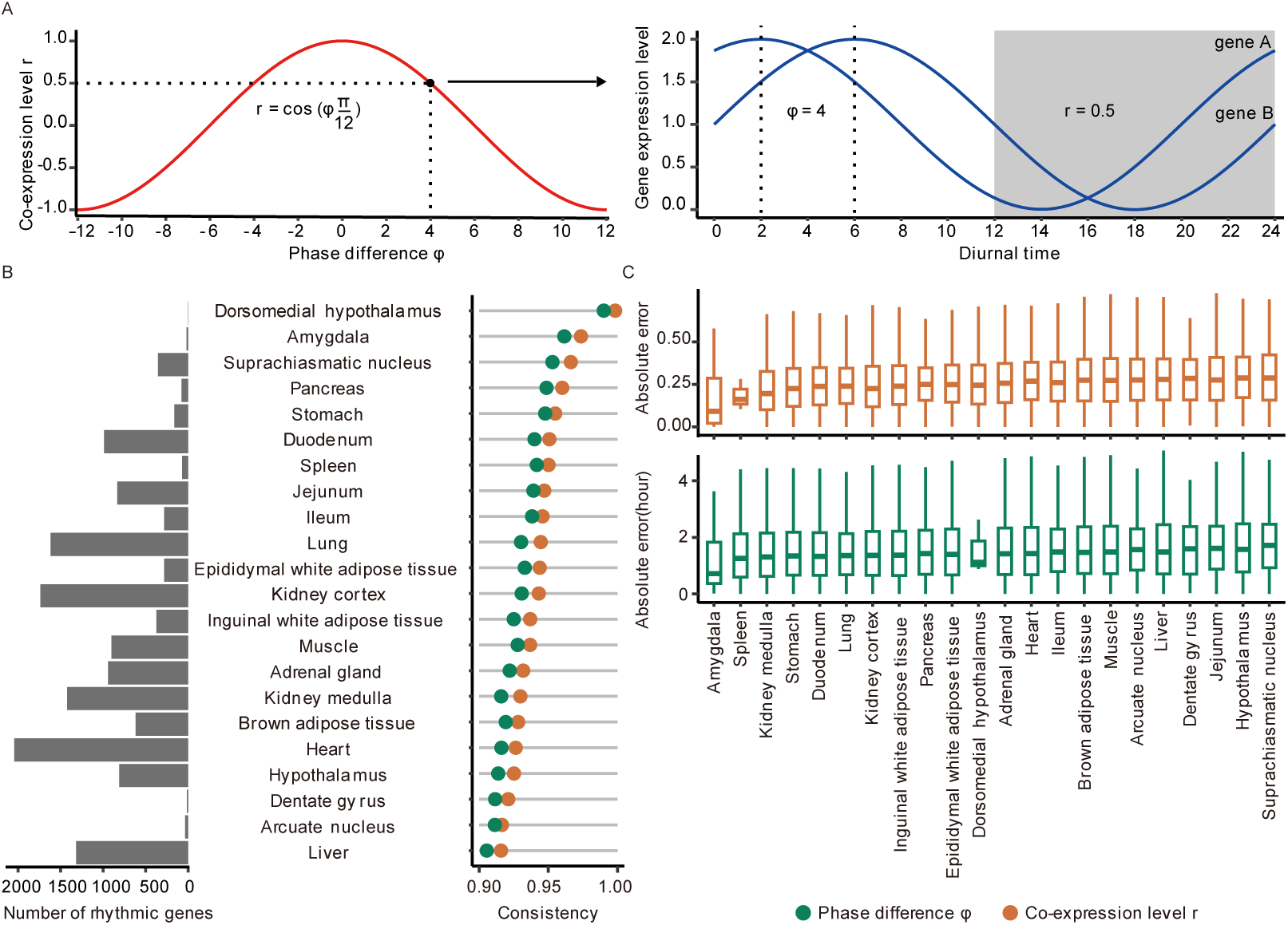
Phase differences can largely account for the co-expression levels between rhythmic genes. (A) The co-expression level r (red) between rhythmic genes (blue) described by sine functions are equal to the cosine of the phase difference φ. (B) The number of rhythmic genes, the consistency between observed and predicted co-expression levels, and the consistency between observed and predicted phase differences across different tissues in mice. (C) The absolute error between observed and predicted co-expression levels of rhythmic genes, and the absolute error between observed and predicted phase differences of rhythmic genes across different tissues in mice.

The results demonstrated that across 22 mouse tissues, the consistency between observed and predicted values for both phase differences and co-expression levels exceeded 0.9 (Figure 1B). Excluding the dorsomedial nucleus, which was removed due to having only three rhythmic genes, the amygdala had the highest consistency, with both values exceeding 0.96. Even in the liver, which had the lowest consistency, both values remained above 0.90. The suprachiasmatic nucleus (SCN), the central pacemaker of the circadian system, exhibited the second-highest consistency among all tissues, with both values exceeding 0.95 (Figure 1B). In terms of deviation between predicted and observed values, the smallest mean absolute error for co-expression levels was 0.15 in the amygdala, while the largest was 0.3 in the arcuate nucleus, with a median of 0.265 across all tissues (Figure 1C). The smallest mean absolute error for phase differences was 1.2 hours in the amygdala, while the largest was 1.7 hours in the dentate gyrus, with a median of 1.58 hours across all tissues (Figure 1C). These results indicate that phase differences account for most of the co-expression levels between rhythmic genes, making the latter a reliable proxy for the former.

### The strength of rhythmic expression affects the power of phase differences to explain co-expression levels

The explanation of co-expression levels by phase differences relies on how well the expression curve fits a simple sine function. In practical applications, the p-value for detecting rhythmic expression significance is often used to characterize the strength of rhythmic expression, which is generally related to how well the expression curve fits a sine function^29,33,34^. Thus, we examined how the strength of rhythmic expression affects the power of phase differences to explain co-expression levels. To this end, using mouse diurnal transcriptomic dataset, we ranked genes in each tissue based on the p-values provided by MetaCycle^33^. The top 5000 genes were grouped into sets of 100, and within each group, we calculated the consistency between observed and predicted values as well as the mean absolute error (Figure 2A and 2B).

**Figure 2.**
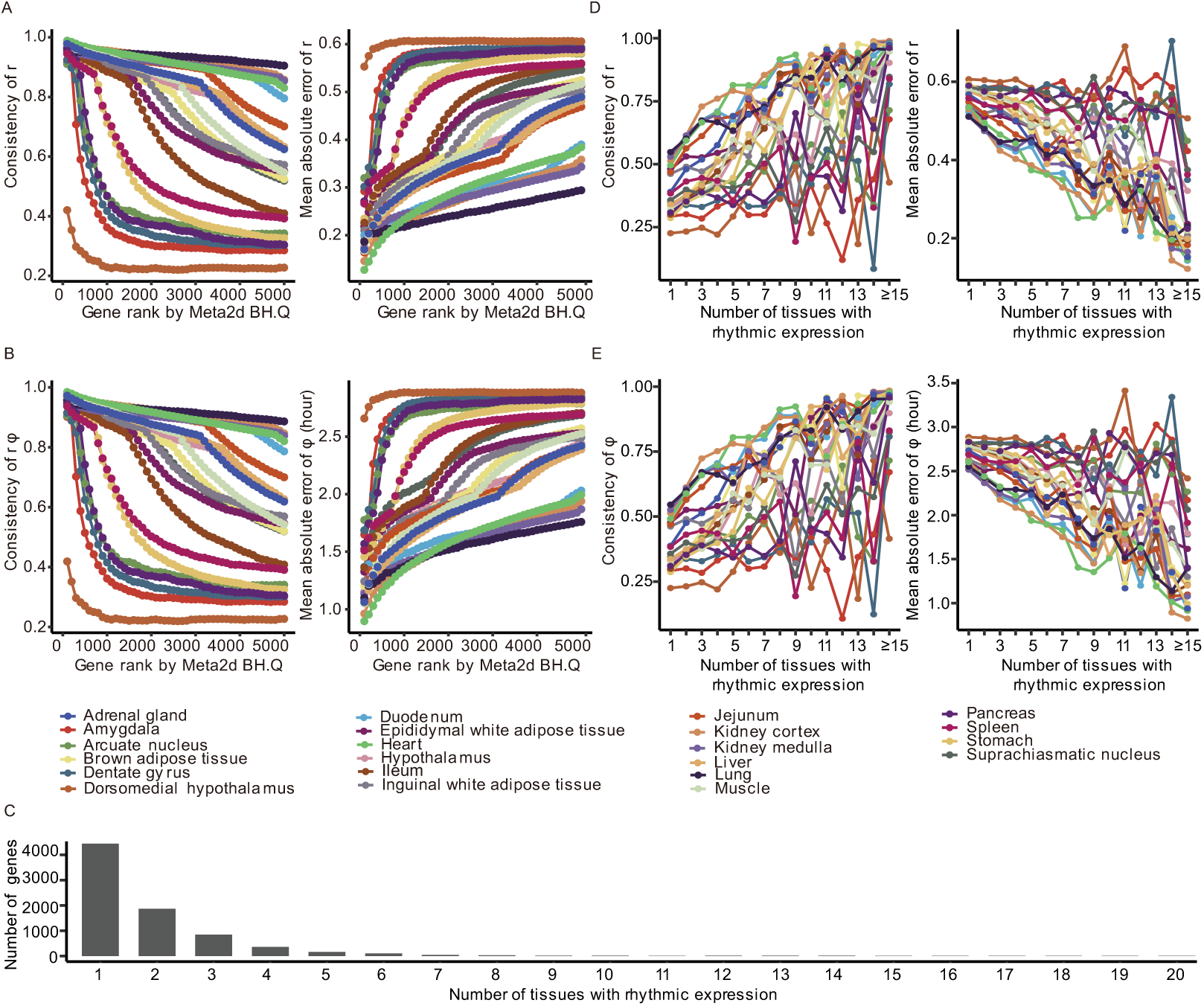
The strength and tissue-specificity of rhythmic expression affect the extent to which phase differences explain co-expression levels. (A) As the strength of rhythmic expression decreases, the consistency between observed and predicted co-expression levels decreases, and the mean absolute error increases across different tissues in mice. (B) As the strength of rhythmic expression decreases, the consistency between observed and predicted phase difference decreases, and the mean absolute error increases across different tissues in mice. (C) The distribution of the number of tissues showing rhythmic expression for rhythmic genes. (D) As the number of tissues with rhythmic expression increases, the consistency between observed and predicted co-expression levels increases, and the mean absolute error decreases across different tissues in mice. (E) As the number of tissues with rhythmic expression increases, the consistency between observed and predicted phase difference increases, and the mean absolute error decreases across different tissues in mice.

The results showed that in all tissues, as the p-value representing the strength of rhythmic expression increased, the consistency between observed and predicted co-expression levels gradually decreased, while the mean absolute error gradually increased (Figure 2A). Similarly, as the p-value increased, the consistency between observed and predicted phase differences also gradually decreased, while the mean absolute error increased (Figure 2B). Notably, in certain tissues, the decline in consistency was relatively gradual; even for genes ranked between 4900 and 5000, the co-expression level consistency remained above 0.80, while the phase difference consistency exceeded 0.79. These tissues included the lung, renal cortex, renal medulla, heart, and duodenum, where the number of rhythmic genes was 1617, 1738, 1422, 2045, and 988, respectively. Moreover, in all tissues except the dorsal nucleus, the number of genes with co-expression consistency above 0.80 was several times greater than the number of rhythmic genes (Figure 2A and 2B). This suggests that in these tissues, even genes not classified as rhythmically expressed still exhibited co-expression patterns largely governed by phase differences. These findings highlight that the strength of rhythmic expression plays a crucial role in determining how well phase differences explain co-expression levels, and this explanatory mechanism may extend beyond rhythmic genes.

### The tissue specificity of rhythmic expression affects the power of phase differences to explain co-expression levels

The circadian rhythmic expression of genes exhibits tissue specificity. Thus, we examined how the tissue specificity of rhythmic expression affects the explanatory power of phase differences for co-expression levels. Specifically, in the diurnal transcriptomic data of 22 mouse tissues, 7906 genes exhibited rhythmic expression in at least one tissue. Among them, 4445 genes (56%) were rhythmically expressed in only one tissue, while only 47 genes showed rhythmic expression in more than 11 tissues (Figure 2C). Next, we grouped genes based on the number of tissues in which they exhibited rhythmic expression and calculated the consistency between observed and predicted values as well as the mean absolute error within each group (Figure 2D and 2E).

The results showed that although fluctuations existed in some tissues, overall, as the number of tissues with rhythmic expression increased, the consistency between observed and predicted co-expression levels gradually increased, while the mean absolute error gradually decreased (Figure 2D and 2E). Similarly, as the number of tissues with rhythmic expression increased, the consistency between observed and predicted phase differences also gradually increased, while the mean absolute error decreased. Importantly, for genes with rhythmic expression in at least 15 tissues, their co-expression consistency exceeded 0.8 in 19 tissues and surpassed 0.9 in 16 tissues (Figure 2D and 2E). Similarly, the consistency of phase differences for these genes exceeded 0.8 in 19 tissues and surpassed 0.9 in 15 tissues (Figure 2D and 2E). In conclusion, the conversion between phase differences and co-expression levels performs more robustly in genes with broad rhythmic expression across multiple tissues.

### Screening of reference genes and construction of tissue-specific reference matrices

Existing methods that use co-expression levels as a proxy for phase differences to assess circadian rhythm disruptions lack systematic selection and validation of reference genes, instead relying on manually selected classical clock genes^20,28^. Since the strong explanatory power of phase differences for co-expression levels is fundamental to these methods, we propose that reference genes should be selected based on strong rhythmic expression across multiple tissues, as supported by the above analysis. Furthermore, as the lists of rhythmic genes in human tissues are derived from predictive algorithms rather than time-series experiments, reference genes should exhibit conserved rhythmic expression in both humans and mice to enhance reliability^21,32,35^.

To this end, we first identified genes with rhythmic expression in more than half of the tissues using multi-tissue transcriptomic data from humans and mice^21,32^. We then clustered all tissues based on the presence or absence of rhythmic gene expression and observed that brain and peripheral tissues segregated into two clusters (Figure S1A). Next, we took the intersection of rhythmic genes between the two species and found that NR1D1 was the only gene retained in brain tissues. Therefore, we focused on peripheral tissues in subsequent analyses. In peripheral tissues, 13 genes were found to be conserved between species: *ADAMTS4*, *ARNTL*, *BHLHE41*, *CIART*, *DBP*, *HLF*, *NFIL3*, *NPAS2*, *NR1D1*, *NR1D2*, *PER2, PER3*, and TEF (Figure S1B). This set of genes contains major clock genes, but notable classical clock genes such as *CRY1*, *CRY2*, and *PER1* were absent. Notably, *ADAMTS4*, also known as aggrecanase-1, plays a crucial role in cartilage degradation by cleaving aggrecan, making it a key target in osteoarthritis research^36,37^. Although previous studies have confirmed the circadian rhythmic expression of *ADAMTS4* in chondrocytes, its widespread rhythmic expression across multiple tissues remains an area of interest for further investigation^38,39^.

Next, we specifically examined the conversion between co-expression levels and phase differences for these 13 genes. To achieve this, we calculated the consistency and absolute error between observed and predicted values for these 13 genes using multi-tissue transcriptomic data from mice and humans (Figure S2). The results showed that for both co-expression levels and phase differences, the consistency of these genes exceeded 0.9 in peripheral tissues of mice, while in humans, 20 tissues exhibited consistency above 0.9 (Figure S2A). Furthermore, in peripheral tissues of mice, the mean absolute error of co-expression levels ranged from 0.12 to 0.26, with a median of 0.19, while the mean absolute error of phase differences ranged from 0.90 to 1.7 hours, with a median of 1.4 hours. In humans, the mean absolute error of co-expression levels across tissues ranged from 0.42 to 0.62, with a median of 0.49, while the mean absolute error of phase differences ranged from 2.4 to 3.5 hours, with a median of 2.8 hours (Figure S2B). Considering these findings, we selected the 13 genes that exhibit widespread rhythmic expression in peripheral tissues of both humans and mice as reference genes for constructing tissue-specific reference matrices.

Thus, for mice, we utilized multi-tissue transcriptomic data under *ad libitum* feeding conditions^32^. For each tissue, we extracted the expression values of the reference genes, calculated their co-expression levels, and converted them into phase differences, ultimately constructing tissue-specific reference phase difference matrices (Figure S3). For humans, we used data from the GTEx project and applied the same approach to derive reference phase difference matrices for each tissue (Figure S4)^21^. It can be observed that the patterns of these phase difference matrices exhibit a high degree of conservation across species and tissues.

### Construction of the phase difference variation inference method

Existing methods that evaluate circadian rhythm disruptions based on co-expression levels lack analyses specifically addressing phase difference variation in individual gene pairs^20,28^. To address this, we refined the existing methodology by developing a phase difference variation (Δφ)-based approach, leveraging the previously established tissue-specific reference matrices to detect gene pairs exhibiting significant phase difference alterations. Briefly, we calculated the phase difference matrix for the input data, subtracted the reference matrix from it to obtain the phase difference variation matrix, and used the distribution of phase difference variations across all expressed genes as the null distribution to derive p-values (Figure 3).

**Figure 3.**
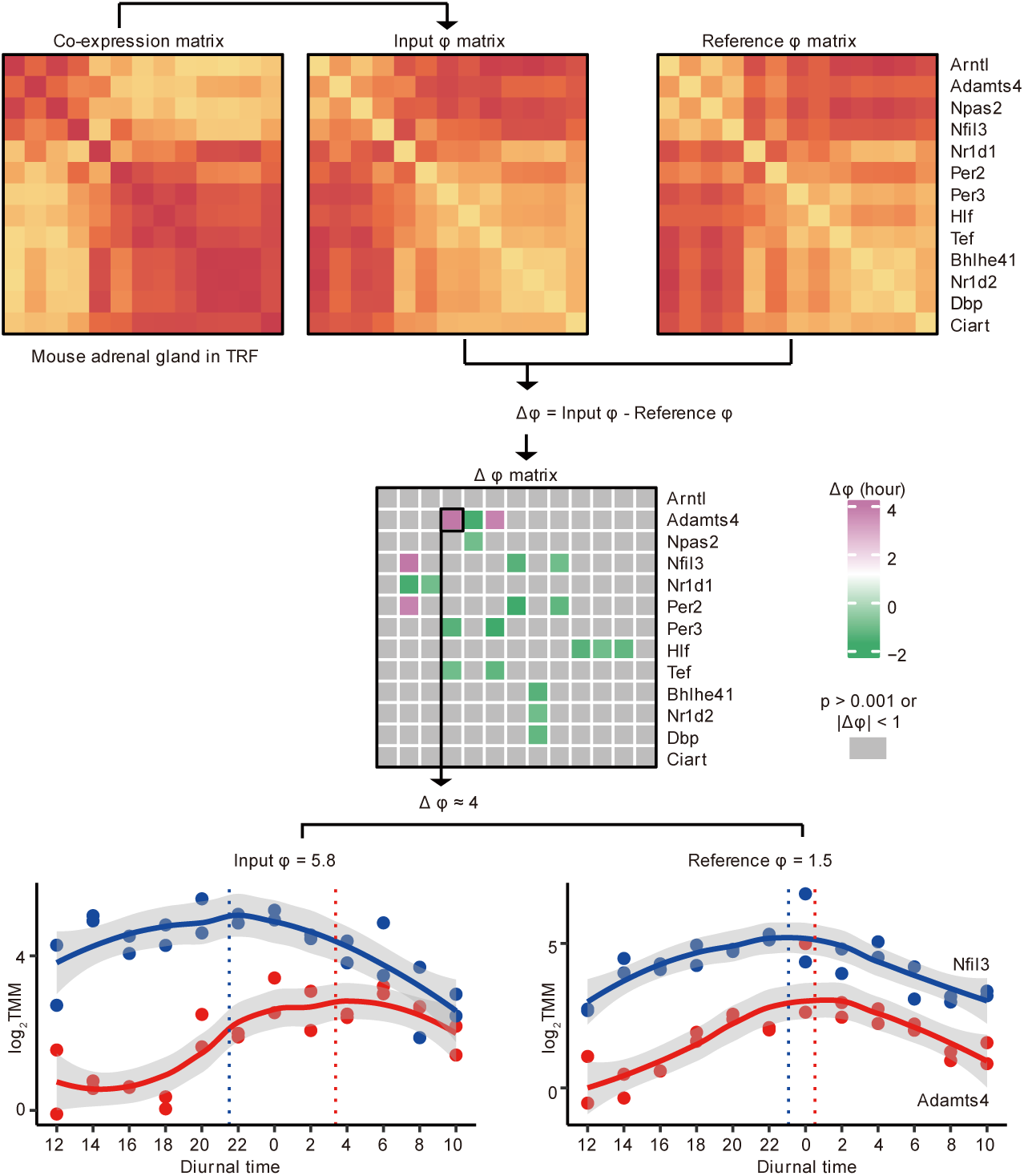
The workflow of the phase difference variation inference method. We use transcriptomic data from the adrenal gland of mice under time-restricted feeding conditions as an example to demonstrate the workflow of this method. First, we extracted the expression values of reference genes, computed the co-expression matrix between reference genes, and converted it into a phase difference matrix. Then, we subtracted the reference matrix of the corresponding tissue from this matrix to obtain the phase difference variation matrix, and computed the significance of each phase difference variation. Finally, we applied a threshold where the absolute phase difference variation exceeded 1 hour and the p-value was less than 0.001 to identify gene pairs with significant phase difference changes in the input data. The results showed that the largest phase difference variation in this tissue was 4 hours between Adamts4 and Nfil3. Consistent with expectations, in the real time-series data, the phase differences between Adamts4 and Nfil3 under the two feeding conditions were 1.5 hours and 5.8 hours, respectively, with a variation of 4.3 hours. The inferred 4 hours was very close to this.

To demonstrate the workflow of this method, we take transcriptomic data from the adrenal gland of mice under time-restricted feeding conditions as an example (Figure 3). First, we extracted the expression values of the reference genes, calculated the co-expression matrix among them, and converted it into a phase difference matrix. Next, we subtracted the reference matrix of the corresponding tissue from this matrix to obtain the phase difference variation matrix, while also calculating the statistical significance of each phase difference variation. Finally, by applying a threshold where the absolute value of the phase difference variation exceeds 1 hour and the p-value is less than 0.001, we identified gene pairs with significant phase difference variations in the input data. The results indicated that the greatest phase difference variation in this tissue was 4 hours between *Adamts4* and *Nfil3*. As expected, in the actual time-series data, the phase differences between *Adamts4* and *Nfil3* under the two feeding conditions were 1.5 hours and 5.8 hours, respectively, yielding a variation of 4.3 hours, which closely matches the inferred value of 4 hours (Figure 3).

Subsequently, to comprehensively evaluate the effectiveness of the proposed method, we applied it to the diurnal transcriptomic data of mouse peripheral tissues under time-restricted feeding conditions and compared the calculated significant phase difference variations with observed values (Figure S5A). The results showed that the consistency of phase difference variations in these tissues ranged from 0.09 to 1.0, with a median of 0.77, while the mean absolute error ranged from 0.47 to 1.76 hours, with a median of 0.91 hours (Figure S5A). These results demonstrate that the method has good predictive performance for phase difference variations. Lastly, to assess the global alterations in circadian regulatory networks under varying conditions, we employed the Euclidean distance between phase difference matrices as a metric. Using the mouse time-restricted feeding dataset as an example, we filtered out non-significant gene pairs and then calculated the Euclidean distance between phase difference matrices under the two conditions (Figure S5B). The results showed that in epididymal white adipose tissue, the phase differences between reference genes, most of which are clock genes, were strongly affected by time-restricted feeding, whereas the lung was the least affected tissue (Figure S5B). Additionally, by visualizing all significantly altered gene pairs, we observed that the phase difference of the same gene pair exhibited tissue-specific directional changes. For instance, the phase difference between *Adamts4* and *Nfil3* significantly increased in epididymal white adipose tissue, adrenal gland, inguinal white adipose tissue, muscle, and liver, but significantly decreased in the renal cortex (Figure S5C). These results suggest that time-restricted feeding influences clock gene expression in a tissue-specific manner.

### Widespread alterations in phase differences between clock genes in cancer

To investigate the applicability of the phase difference variation inference method, we conducted a protein-protein interaction (PPI) analysis on the 13 reference genes to uncover the most relevant biological processes^40^. The results showed that genes directly interacting with the reference genes, which are predominantly clock genes, were enriched in pathways related to the Wnt signaling pathway, cell cycle, microRNAs in cancer, and transcriptional dysregulation in cancer, indicating a strong association between clock genes and cancer (Figure S6).

Thus, in the TCGA cancer transcriptomic data, we retained eight cancer types for which corresponding tissue-specific reference phase difference matrices from GTEx were available: lung adenocarcinoma (LUAD), lung squamous cell carcinoma (LUSC), stomach adenocarcinoma (STAD), thyroid carcinoma (THCA), breast invasive carcinoma (BRCA), prostate adenocarcinoma (PRAD), uterine corpus endometrial carcinoma (UCEC), and colon adenocarcinoma (COAD). For each cancer type, we input the tumor and normal sample data into the phase difference variation inference method, using a threshold where the absolute phase difference variation exceeded 2 hours and the p-value was below 0.001, to identify gene pairs with significant phase difference variation in both tumor and normal samples (Figure 4A). The results showed that tumor samples exhibited more significantly altered gene pairs in phase differences than normal samples, with a relatively balanced number of gene pairs showing increased and decreased phase differences. However, a certain number of significantly altered gene pairs were also observed in normal samples, most of which exhibited decreased phase differences, and LUSC did not show any significantly altered gene pairs (Figure 4A). Considering that normal samples in TCGA are largely derived from tumor-adjacent tissues, these findings imply that the rhythmic expression of clock genes in tumor-adjacent tissues may differ from that in healthy tissues.

**Figure 4.**
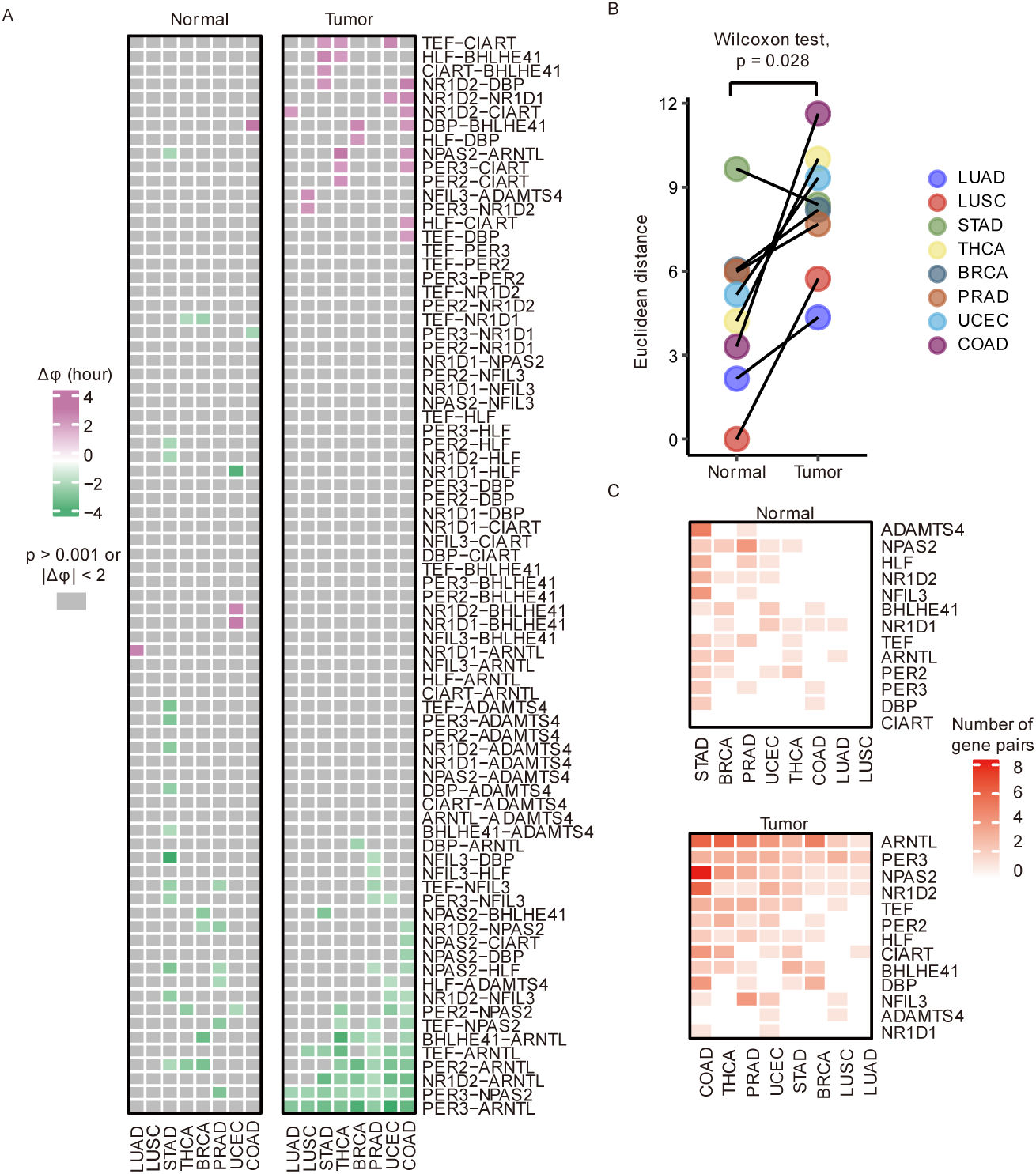
Phase differences among clock genes exhibit widespread alterations in cancer. (A) Gene pairs showing significant phase difference variations relative to the reference in normal and tumor samples from cancer patients. (B) The Euclidean distance between the phase difference matrix of each cancer and the reference matrix. (C) The count of gene pairs showing significant phase difference variations linked to each reference gene.

Next, we quantified the deviation of the circadian regulatory network relative to healthy individuals by calculating the Euclidean distance between the phase difference matrix of these samples and the reference matrix (Figure 4B). The findings revealed that, in general, tumor samples had a significantly larger Euclidean distance (Wilcoxon test, p = 0.028). However, normal samples of STAD exhibited a larger Euclidean distance, with all significant phase difference variation between gene pairs being decreases (Figure 4A and 4B). Tumor samples of COAD had the greatest Euclidean distance and the highest number of significantly altered gene pairs (Figure 4A and 4B). Normal samples of BRCA and PRAD had Euclidean distances comparable to those of tumor samples, which may suggest that the rhythmic expression of clock genes in tumor-adjacent tissues of these cancers is also disrupted (Figure 4B).

Subsequently, we counted the number of gene pairs with significant phase difference variation associated with each reference gene (Figure 4C). The results showed that in tumor samples, *ARNTL*, *PER3*, *NPAS2*, and *NR1D2* were involved in significantly altered gene pairs across all eight cancer types, with ARNTL having the highest number of associated gene pairs, while *NR1D1* and *ADAMTS4* exhibited the fewest changes (Figure 4C). Furthermore, *ADAMTS4* was extensively involved in significantly altered gene pairs only in normal samples of STAD, while it was scarcely involved in STAD and other cancerous tumor samples, which may explain the larger Euclidean distance observed in STAD normal samples (Figure 4B and 4C). In conclusion, by applying the phase difference variation inference method to cancer datasets, we uncovered widespread and cancer-type-specific significant alterations in phase differences between clock genes in tumors.

### The single-sample circadian rhythm score ssDistance stratified cancer patients into subgroups with different survival outcomes

To explore circadian rhythm-related heterogeneity across a large number of samples, we developed a single-sample circadian rhythm score based on phase differences. Specifically, for each iteration, we excluded one input sample, constructed the phase difference matrix using the remaining samples, calculated its Euclidean distance from the reference matrix, and normalized the result to obtain the single-sample distance (ssDistance) for the excluded sample. Hence, a higher ssDistance score suggests that the phase differences among clock genes in the sample show less deviation from the reference matrix (Figure 5A).

**Figure 5.**
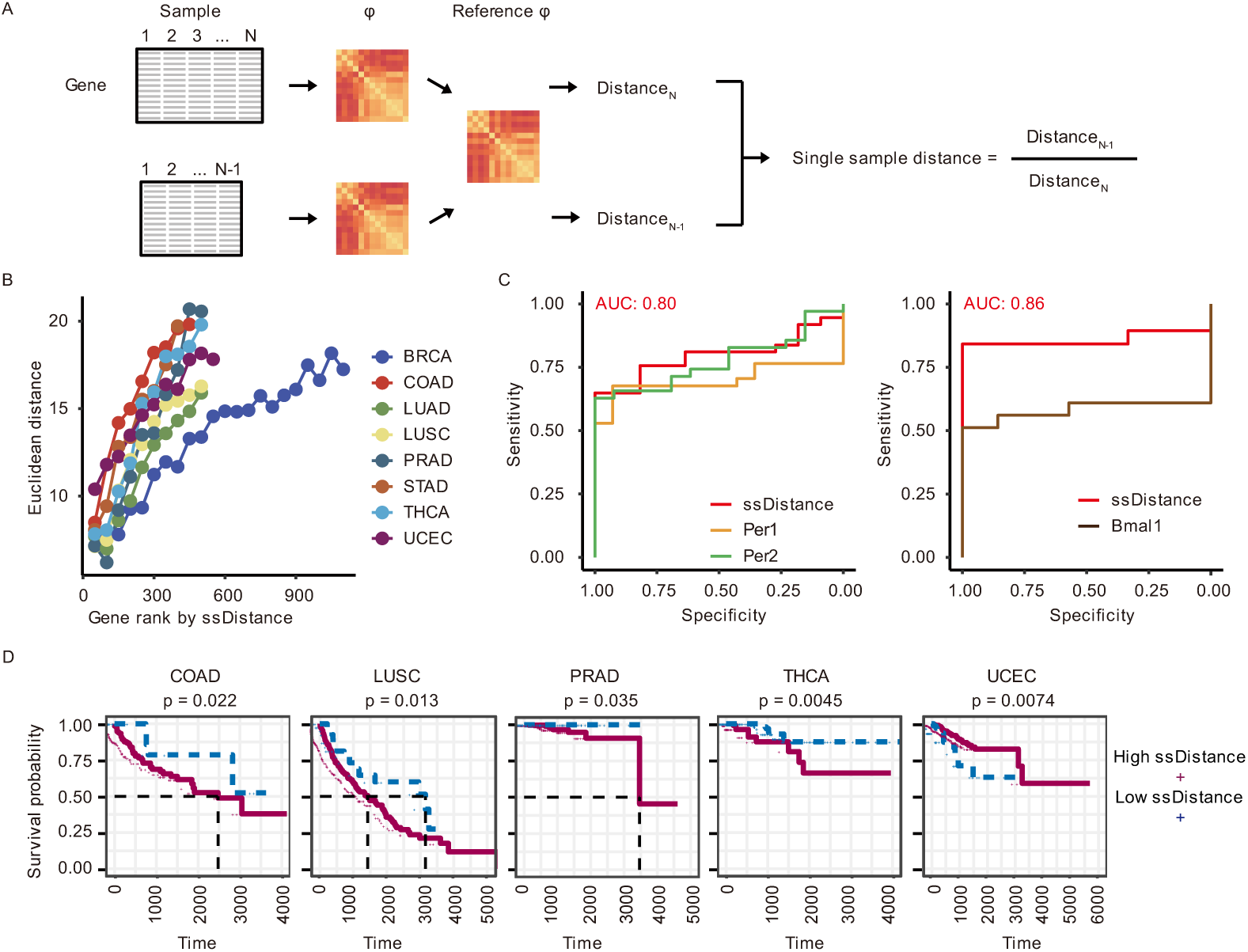
The single-sample circadian rhythm score ssDistance distinguished different overall survival in cancer patients. (A) The workflow of the phase difference-based single-sample circadian rhythm score ssDistance. (B) As the ssDistance increases, the overall Euclidean distance for each group of 50 samples also increases. (C) The ability of the ssDistance to differentiate clock gene knockout samples from control samples is superior to that of expression values.. (D) ssDistance stratified cancer patients into subgroups with different survival durations..

Next, to validate the internal consistency of ssDistance, we sorted the samples in cancer datasets with available reference matrices based on ssDistance and calculated the Euclidean distance between the phase difference matrix and the reference matrix for every 50 samples. We observed that the Euclidean distance increased with increasing ssDistance, indicating a consistent trend between the two (Figure 5B). Furthermore, to assess whether ssDistance can effectively identify samples with circadian rhythm disruptions, we analyzed data from *Per1*/*2* double knockout and *Bmal1* knockout mouse liver samples (Figure 5C)^41^. Specifically, we used ssDistance and the expression levels of the knocked-out genes as metrics and compared their performance in distinguishing treated and control samples based on the area under the curve (AUC). The results showed that ssDistance achieved an AUC of 0.80 in distinguishing *Per1*/*2* double knockout samples and an AUC of 0.86 in distinguishing *Bmal1* knockout samples, both outperforming the classification accuracy based on the expression levels of the knocked-out genes (Figure 5C).

Thus, we applied ssDistance to the TCGA cancer dataset. For the eight cancers with GTEx reference matrices, we calculated ssDistance for each tumor sample, stratified patients into high and low ssDistance groups based on the optimal cutoff, and performed Kaplan-Meier survival analysis for the subgroups (Figure 5D). We observed that in five cancer types, patients were stratified into subgroups with significantly different overall survival. Among them, patients with high ssDistance in COAD (log-rank test, p = 0.022), LUSC (log-rank test, p = 0.013), PRAD (log-rank test, p = 0.035), and THCA (log-rank test, p = 0.0045) had significantly shorter overall survival, suggesting that in these four cancers, patients whose tumor clock gene phase differences deviated more from the reference matrix had longer overall survival. High ssDistance was associated with significantly longer overall survival only in UCEC log-rank test, (p = 0.0074) (Figure 5D).To further validate the association between ssDistance and overall survival across different cancer types, we replaced GTEx samples with TCGA normal samples to generate the reference phase difference matrices and reassessed ssDistance scores in 14 cancers with sufficient sample sizes (Figure S7). We observed that in seven cancer types, patients were stratified into subgroups with significantly different overall survival. Specifically, in bladder carcinoma (BLCA) (log-rank test, p = 0.00053), COAD (log-rank test, p = 0.0056), head and neck squamous cell carcinoma (HNSC) (log-rank test, p = 0.0015), and LUSC (log-rank test, p = 0.03), patients with high ssDistance had significantly shorter overall survival. In kidney renal clear cell carcinoma (KIRC) (log-rank test, p < 0.0001), kidney renal papillary cell carcinoma (KIRP) (log-rank test, p = 0.011), and UCEC (log-rank test, p = 0.015), patients with high ssDistance had significantly longer overall survival. Notably, the findings for COAD, LUSC, and UCEC were consistent with the results obtained using healthy samples as the reference (Figure S7). These findings suggest that the single-sample circadian rhythm score ssDistance, based on phase differences between clock genes, plays a dual role—potentially preventing certain cancers while increasing the risk of others—highlighting its cancer-type-specific nature.

## Discussion

In this study, we examined and refined the method for inferring phase differences from clock gene co-expression in timestamp-free samples. We found that the co-expression levels between rhythmic genes are primarily determined by phase differences, and this relationship is influenced by the strength of rhythmic expression and tissue specificity. Therefore, we selected evolutionarily conserved multi-tissue rhythmic genes in mice and humans and constructed tissue-specific phase difference matrices, upon which we developed a method for inferring phase difference variations. Importantly, we extended this method to the single-sample level by developing a phase difference-based circadian rhythm score, ssDistance. The application of these methods in cancer datasets highlights the cancer-type-specific alterations in clock gene phase differences and the complex effects of clock gene rhythmic expression on patient survival. We hope that these methods will enable researchers to investigate molecular circadian rhythms in extensive public datasets that lack circadian time annotations.

Previous studies have discussed the incorporation of phase difference information in the co-expression of clock genes and validated this in transcriptomic data from mouse lung^28^. Our study expanded this scope to 22 mouse tissues and examined nearly all rhythmic genes, including clock genes, confirming the generality of this mathematical relationship. Furthermore, we found that the co-expression of many genes not traditionally considered rhythmically expressed could also be explained by phase differences. This suggests that even for non-rhythmic genes, the peak expression time has a strong influence on co-expression. It may also indicate that current rhythmic expression detection algorithms underestimate the number of genes whose expression curves fit a simple sine function. On the other hand, previous studies constructed reference co-expression matrices using mouse tissues and described overall differences by computing distances between matrices^20^. Our study, in contrast, used human transcriptomic data to construct tissue-specific phase difference matrices as references and incorporated statistical significance calculations for phase difference variations in individual gene pairs, allowing for more precise identification of significantly affected gene pairs. Lastly, previous studies identified a complex relationship between the expression levels of individual clock genes and cancer risk^16^. In contrast, our study employed a phase difference-based single-sample scoring method to evaluate phase difference—a key circadian rhythm parameter—in the context of survival, highlighting the significance of examining the global properties of circadian regulatory networks in tumors.

This study still presents several unresolved challenges. First, the assumption that the expression curves of reference genes conform to a sine function is a fundamental premise of our method. While we attempted to ensure this by selecting evolutionarily conserved multi-tissue rhythmic genes, there is still a lack of research on the 24-hour time-series expression of these genes *in vivo*, particularly in diseased tissues such as tumors^42–44^. Therefore, this assumption should be taken into account when applying our method. Second, while it is well established that the human circadian cycle remains highly stable under environmental synchronization, mutations in core clock genes can still lead to shortened or extended circadian periods^9,45–48^. Since changes in cycle length can also influence the co-expression of clock genes, our current method struggles to distinguish between the effects of cycle length and phase differences. Despite these limitations, our study provides a powerful tool for investigating circadian rhythms in large-scale public datasets.

## Methods

### Data download and preprocessing

The diurnal transcriptomic data of 22 mouse tissues under two feeding conditions were obtained from the study published by Deota et al^32^. They subjected two groups of mice to either *ad libitum* feeding or time-restricted feeding and collected tissue samples from 22 organs every 2 hours over a 24-hour period for gene expression profiling. They performed TMM normalization on sequencing counts. The MetaCycle R package was used to identify rhythmic genes, with a detection threshold set at a meta2d_BH.Q value of less than 0.05. We downloaded the gene expression data for all tissues under both feeding conditions, along with the corresponding rhythmic gene detection results, from the supplementary materials of the study.

The human multi-tissue transcriptomic data and sample circadian time information were obtained from the study published by Talamanca et al^21^. They integrated data from the GTEx project with the CHIRAL algorithm, assigning an inferred circadian time to each sample. The counts data were then normalized for library size, scaled using the TMM method in edgeR, converted to CPM, and subsequently log-transformed. They performed sine regression on gene expression across all tissues and identified rhythmic genes based on a Benjamini-Hochberg corrected p-value threshold of less than 0.2 and an amplitude greater than 0.5. We directly downloaded the CPM expression values for all samples, the circadian time information for each sample, and the rhythmic gene detection results from the supplementary materials of the study.

TCGA data were downloaded using the GDCquery function from the R package TCGAbiolinks, obtaining gene expression TPM and counts for tumor and control samples across all 33 cancer types in the TCGA project, along with corresponding clinical data^49^. For tumor samples, days to death and last follow-up days were combined to define overall survival time for survival analysis.

The transcriptomic data of clock gene knockouts in mice were obtained from the study published by Aviram et al^41^. They collected liver samples from wild-type, *Per1/2* double knockout, and *Bmal1* knockout mice at 4-hour intervals over a 48-hour period. Gene expression counts for this dataset were obtained from Gene Expression Omnibus (GEO) under accession GSE171975 and converted to TPM values using the GTF file from GENCODE vM24^50^.

### Linking co-expression levels with phase differences

The correlation coefficient r between two sine functions equals the cosine of their phase difference φ:

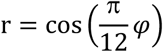

Ideally, the rhythmic expression of clock genes can be accurately described by sine functions. Thus, in the given equation, r denotes the co-expression level between rhythmic genes, and 𝜑 signifies their phase difference. In this study, we substituted the phase differences obtained from rhythmic expression detection algorithms into the equation and used the calculated results as predicted co-expression levels. Simultaneously, we directly computed Pearson correlations from gene expression values and used the results as observed co-expression levels. Similarly, we applied the same approach to compute both the predicted and observed values of phase differences. Subsequently, we assessed the performance of this equation in real data by calculating the consistency and absolute error between predicted and observed values for all gene pairs.

### Screening reference genes and constructing reference matrices

Specifically, in the multi-tissue transcriptomic data of humans and mice, we assigned values of 1 and 0 to rhythmic and non-rhythmic genes, respectively, and selected genes with rhythmic expression in more than half of the tissues. Next, we clustered all tissues based on these genes and found that brain and peripheral tissues formed two distinct clusters. Therefore, we identified the intersection of rhythmic genes in brain and peripheral tissues separately for both species and used them as reference genes. Subsequently, using multi-tissue diurnal transcriptomic data from mice under *ad libitum* feeding conditions, we extracted reference gene expression values for each tissue, calculated co-expression levels among reference genes, and applied the equation to generate phase difference matrices as reference matrices. The same procedure was applied to the human GTEx data to obtain human tissue-specific reference matrices.

### Calculating phase difference variations

To identify gene pairs with significant phase difference variations, we developed a pipeline for calculating phase difference variations (Δφ) and assessing their statistical significance. For a given input dataset, we initially extracted the expression values of reference genes, determined their co-expression levels, and converted them into a phase difference matrix. This matrix was then subtracted from the reference matrix to derive the phase difference variation matrix. To determine the statistical significance of phase difference variations—whether a specific change exceeds the overall background variation—we calculated phase difference variations for all gene pairs in the input dataset and used this distribution as the null distribution to derive p-values. Finally, by applying thresholds on phase difference variations and p-values, we identified gene pairs with significant phase difference variations in the input dataset.

### Calculating single-sample circadian rhythm score

We employed a jackknife resampling strategy, sequentially omitting one sample from the input dataset. The phase difference matrix was calculated using the remaining samples, followed by the computation of its Euclidean distance from the reference matrix. The resulting matrix was then normalized across all samples to derive the single-sample distance (ssDistance) for the omitted sample.

### Protein-protein interaction analysis

We performed a protein-protein interaction analysis on the reference genes to identify biological pathways closely associated with circadian rhythms. Specifically, we input the reference genes into the STRING database, selecting humans as the target species, with a minimum interaction score threshold of 0.8, and performed functional enrichment analysis on the interacting proteins^40^.

### Survival analysis

We estimated the optimal cutoff for ssDistance across multiple cancer types using the R package survminer^51^, stratified patients accordingly, and performed Kaplan-Meier survival analysis using the survfit function from the R package survival^52^, assessing the significance of overall survival differences between the two subgroups.

### Statistical analysis and data visualization

Unless otherwise specified, a p-value threshold of <0.05 was used to determine statistical significance in this study. Data visualization was performed using R (v4.2.0) with various packages, including ggplot2^53^, ggsci^54^, patchwork^55^, ComplexHeatmap^56^, eulerr^57^, and ggplotify^58^.

## Data availability

All relevant data and supplementary materials for this study are deposited in figshare and can be accessed at https://doi.org/10.6084/m9.figshare.28234283.

## Code availability

The code of this publication is available at https://github.com/NextExon/ssDistance_Code.

---

**Figure S1.**
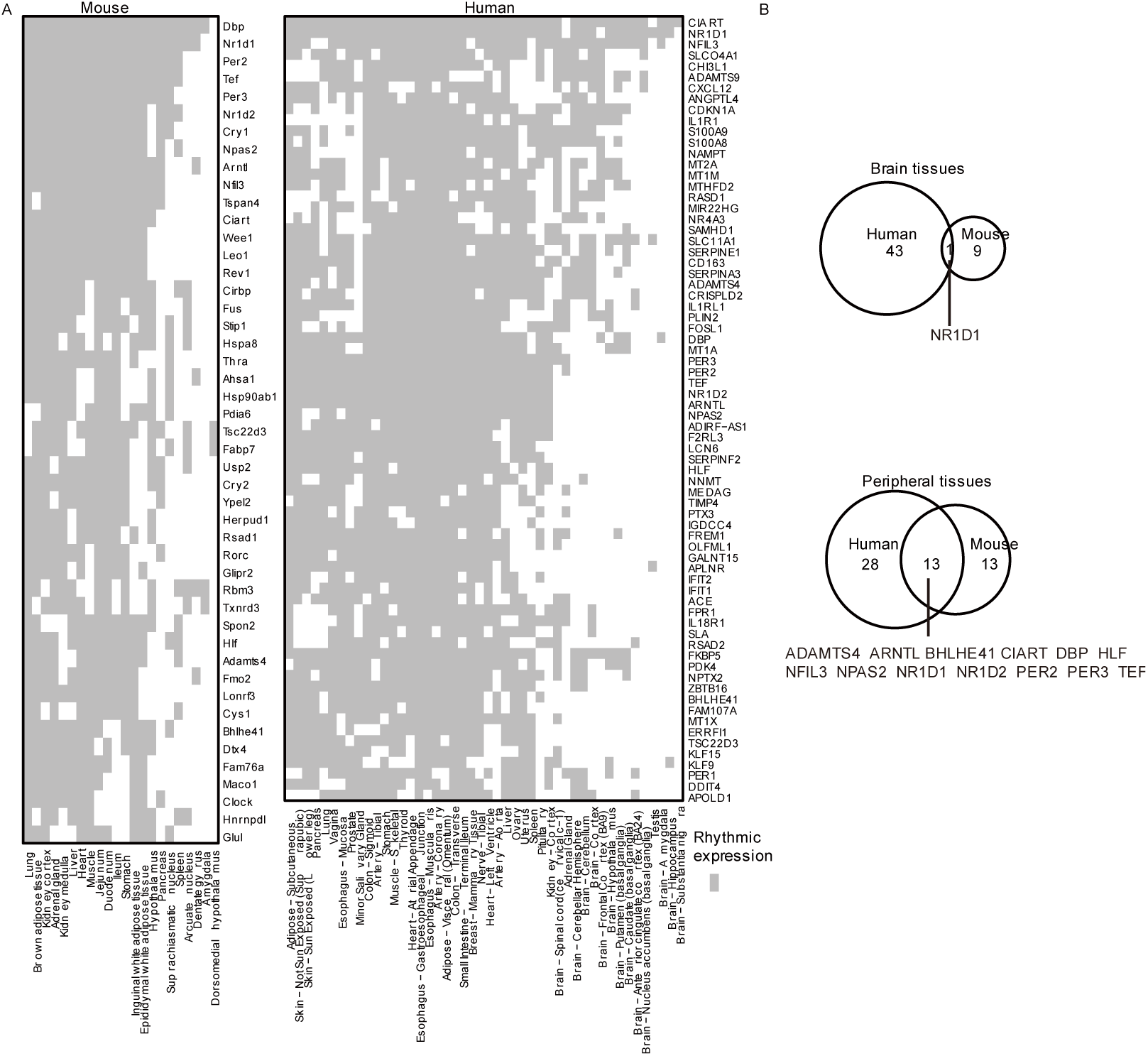
Screening of reference genes. (A) The distribution of genes exhibiting rhythmic expression in over half of the tissues. (B) The intersection of multi-tissue rhythmic genes in humans and mice.

**Figure S2.**
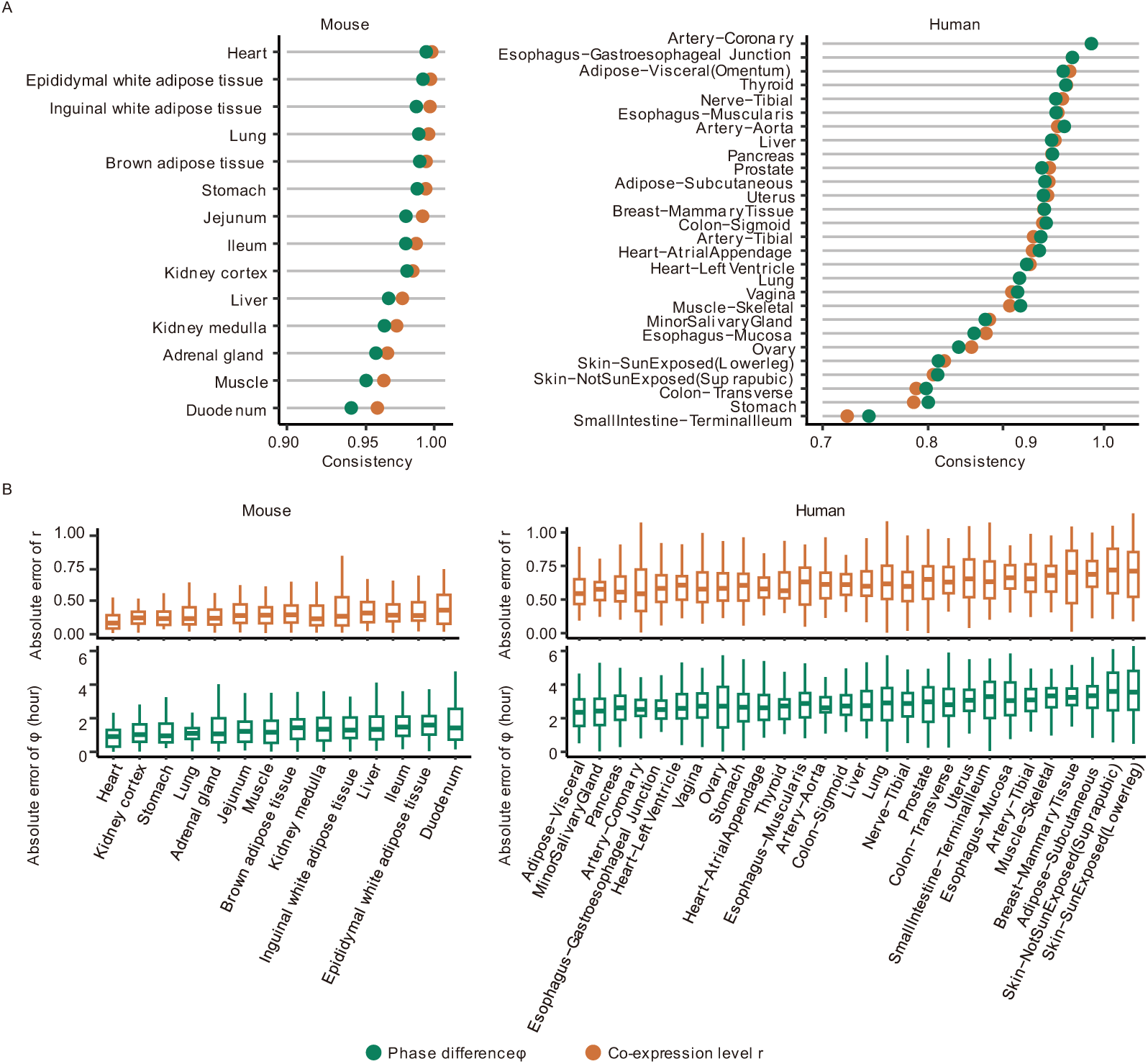
The explanatory power of phase differences between reference genes on co-expression levels. (A) The consistency between observed and predicted co-expression levels, and the consistency between observed and predicted phase differences across different tissues. (B) The absolute error between observed and predicted co-expression levels of rhythmic genes, and the absolute error between observed and predicted phase differences of rhythmic genes across different tissues in mice.

**Figure S3.**
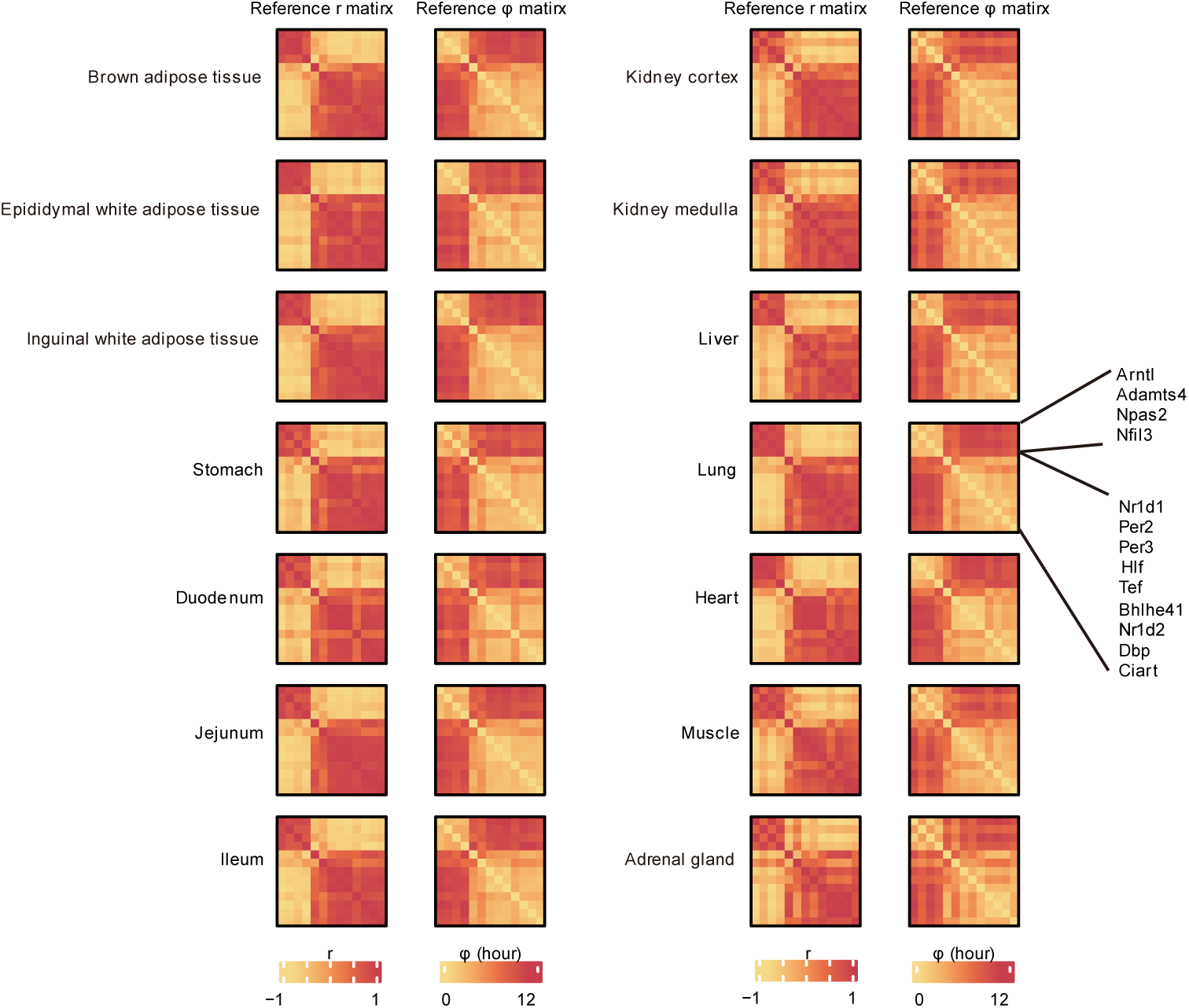
The tissue-specific reference phase difference matrix for mice.

**Figure S4.**
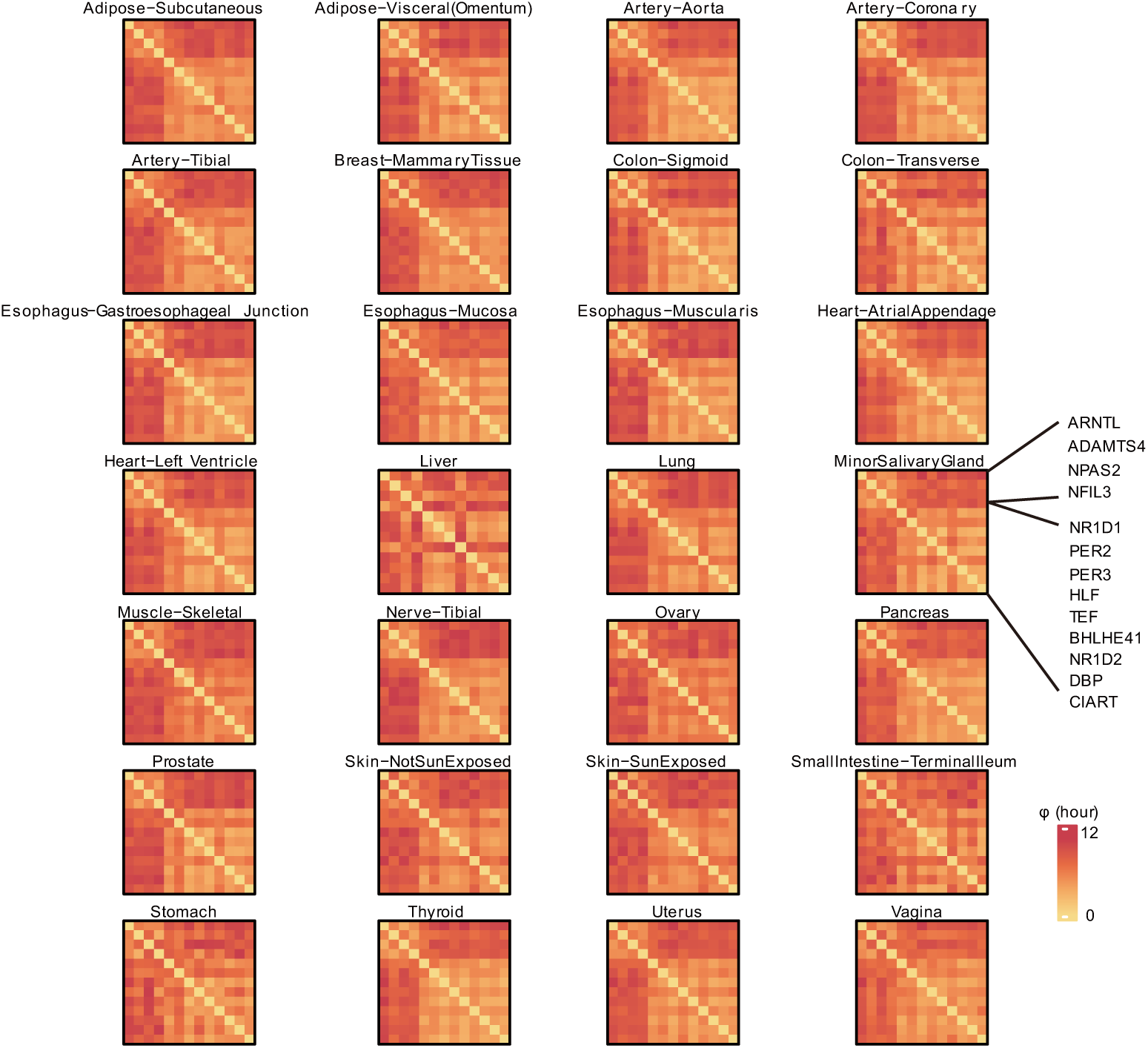
The tissue-specific reference phase difference matrix for humans.

**Figure S5.**
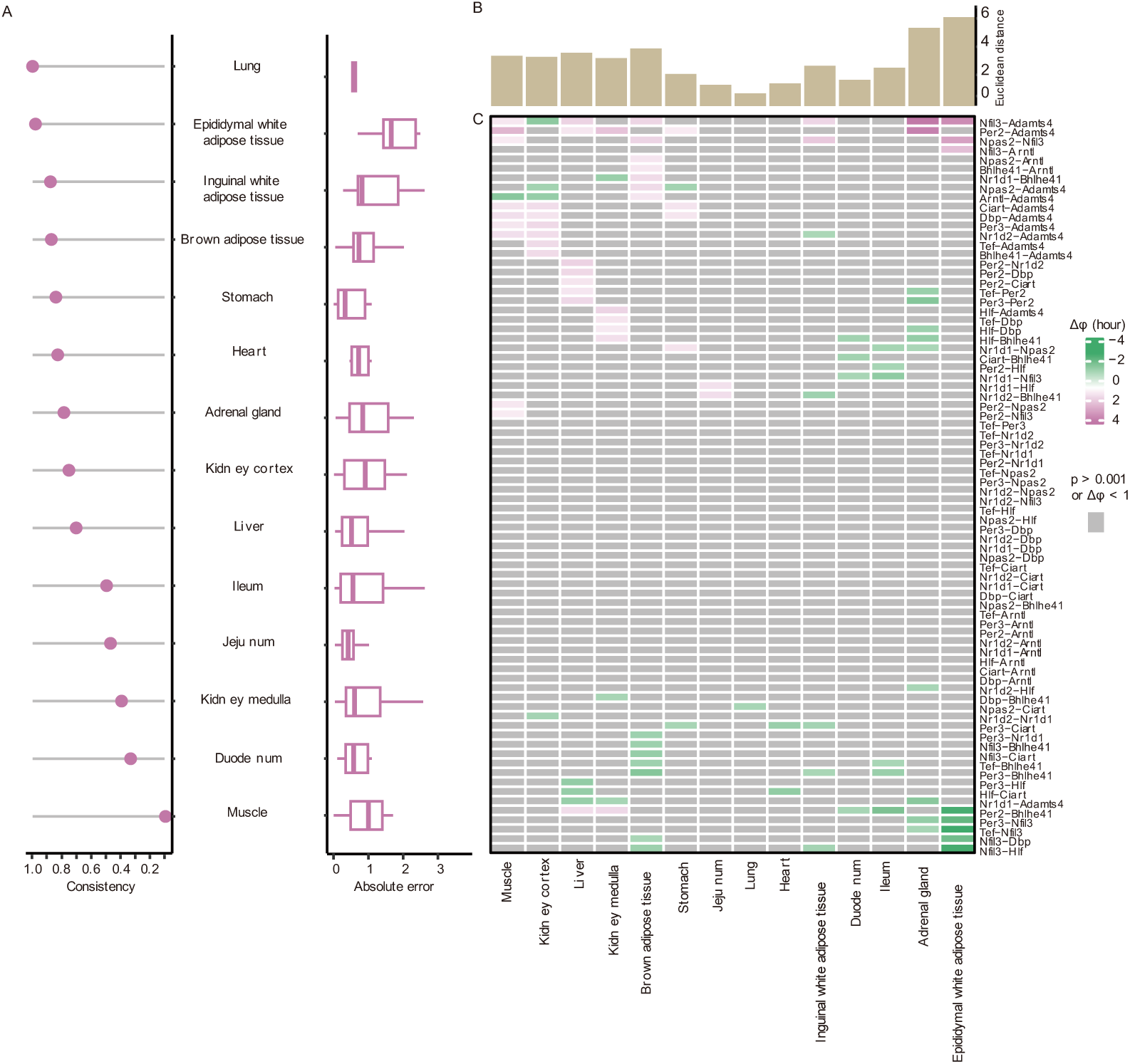
Testing of the phase difference variation inference method. (A) Consistency and absolute error of significant phase difference variations across various mouse tissues. (B) The Euclidean distance between the phase difference matrices of various mouse tissues and the reference matrix under time-restricted feeding conditions. (C) Gene pairs exhibiting significant phase difference variations in different mouse tissues under time-restricted feeding conditions.

**Figure S6.**
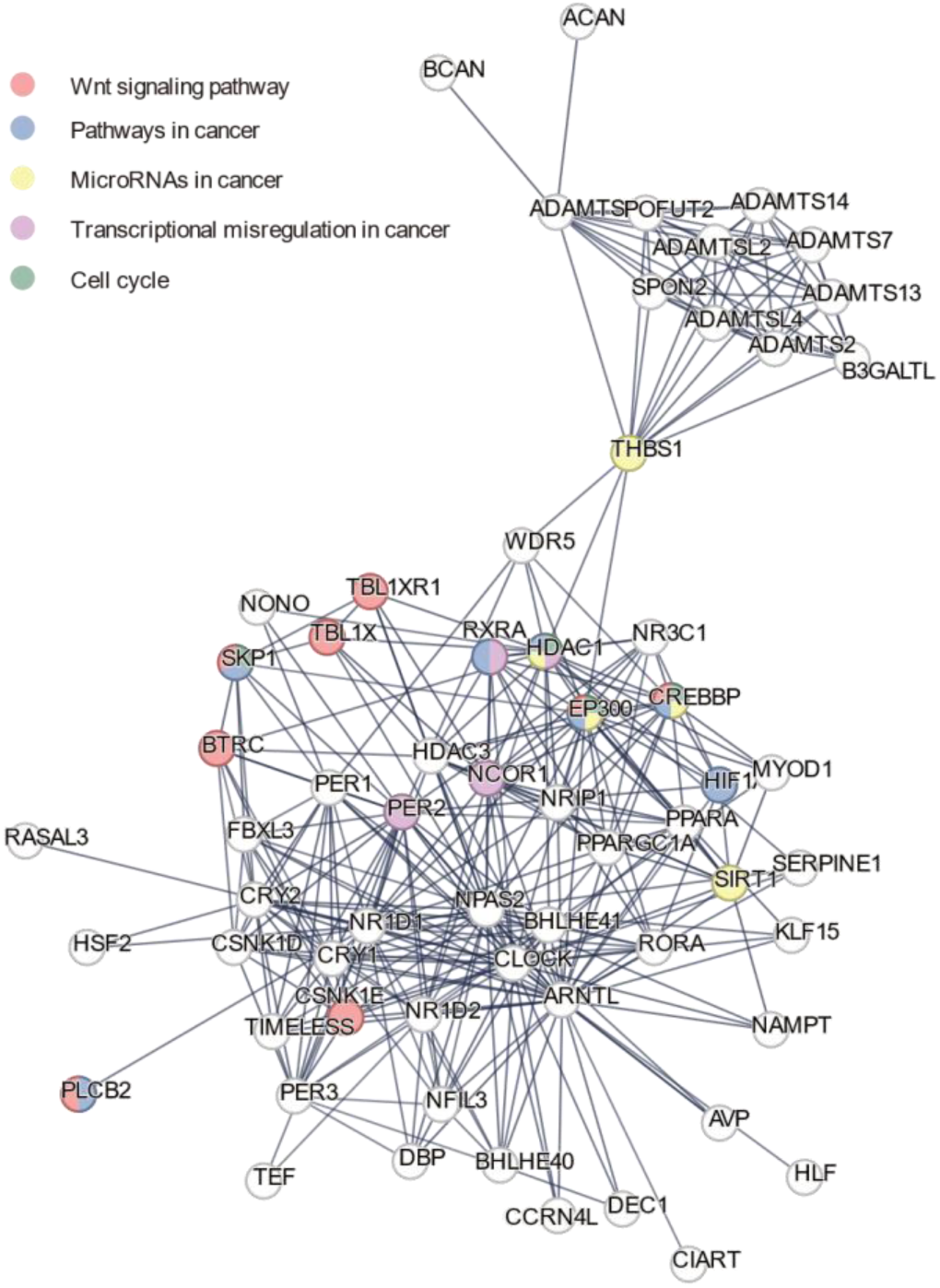
Protein-protein interaction analysis of reference genes.

**Figure S7.**
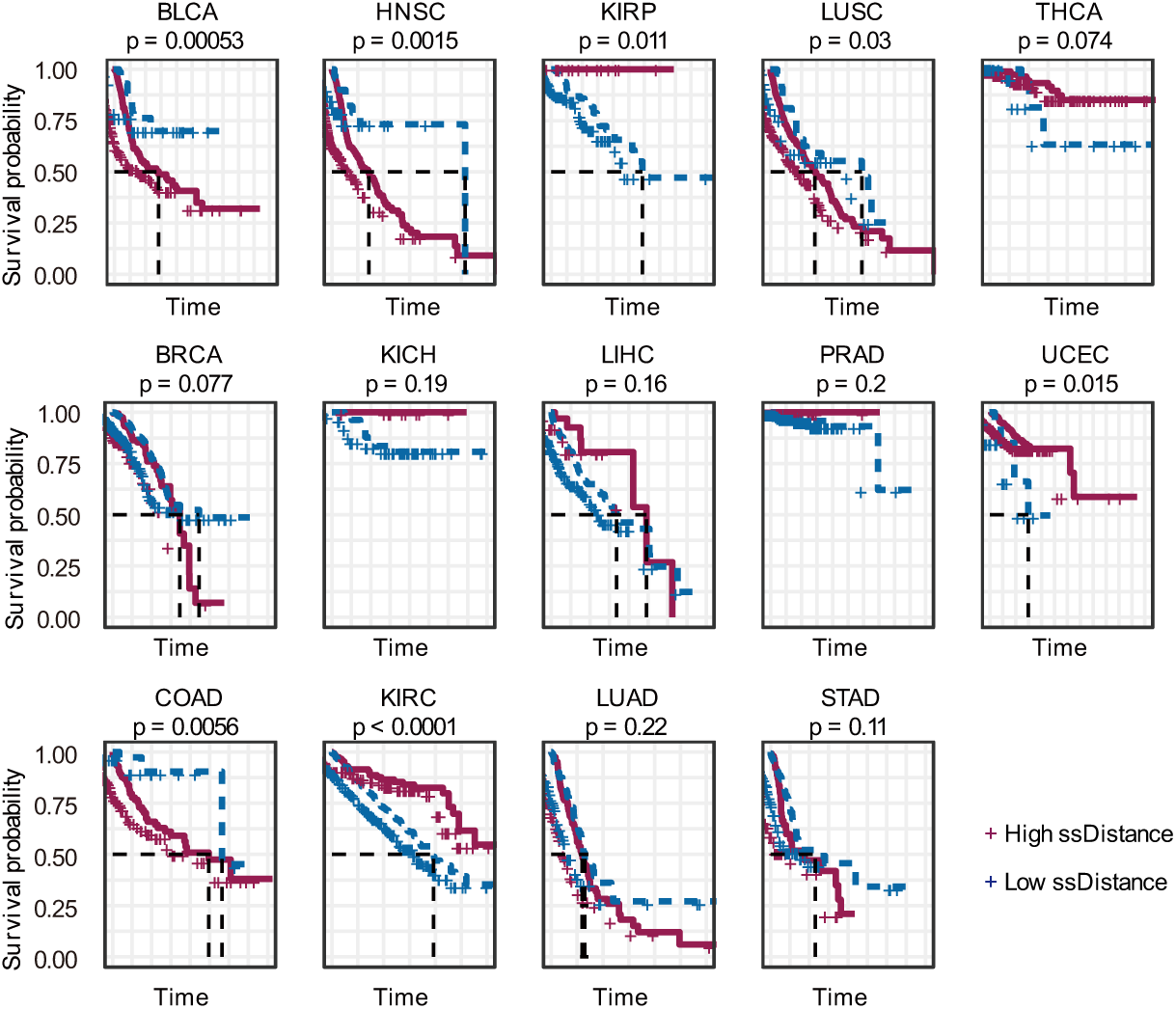
The ssDistance calculated using cancer normal samples as a reference for stratifying survival.

## Reference

1 Paranjpe, D. A. & Sharma, V. K. Evolution of temporal order in living organisms. J Circadian Rhythms 3, 7, doi:10.1186/1740-3391-3-7 (2005).

2 Allada, R. & Bass, J. Circadian Mechanisms in Medicine. N Engl J Med 384, 550–561, doi:10.1056/NEJMra1802337 (2021).

3 Honma, S. The mammalian circadian system: a hierarchical multi-oscillator structure for generating circadian rhythm. J Physiol Sci 68, 207–219, doi:10.1007/s12576-018-0597-5 (2018).

4 Patke, A., Young, M. W. & Axelrod, S. Molecular mechanisms and physiological importance of circadian rhythms. Nat Rev Mol Cell Biol 21, 67–84, doi:10.1038/s41580-019-0179-2 (2020).

5 Rijo-Ferreira, F. & Takahashi, J. S. Genomics of circadian rhythms in health and disease. Genome Med 11, 82, doi:10.1186/s13073-019-0704-0 (2019).

6 Scheiermann, C., Gibbs, J., Ince, L. & Loudon, A. Clocking in to immunity. Nat Rev Immunol 18, 423–437, doi:10.1038/s41577-018-0008-4 (2018).

7 Li, J. Z. et al. Circadian patterns of gene expression in the human brain and disruption in major depressive disorder. Proc Natl Acad Sci U S A 110, 9950–9955, doi:10.1073/pnas.1305814110 (2013).

8 Zhou, L. et al. Circadian rhythms and cancers: the intrinsic links and therapeutic potentials. J Hematol Oncol 15, 21, doi:10.1186/s13045-022-01238-y (2022).

9 Janoski, J. R., Aiello, I., Lundberg, C. W. & Finkielstein, C. V. Circadian clock gene polymorphisms implicated in human pathologies. Trends Genet 40, 834–852, doi:10.1016/j.tig.2024.05.006 (2024).

10 Wang, Z. et al. The interplay of the circadian clock and metabolic tumorigenesis. Trends Cell Biol 34, 742–755, doi:10.1016/j.tcb.2023.11.004 (2024).

11 Kelleher, F. C., Rao, A. & Maguire, A. Circadian molecular clocks and cancer. Cancer Lett **342**, 9–18, doi:10.1016/j.canlet.2013.09.040 (2014).

12 Miro, C. et al. “Time” for obesity-related cancer: The role of the circadian rhythm in cancer pathogenesis and treatment. Semin Cancer Biol 91, 99–109, doi:10.1016/j.semcancer.2023.03.003 (2023).

13 Sancar, A. & Van Gelder, R. N. Clocks, cancer, and chronochemotherapy. Science 371, doi:10.1126/science.abb0738 (2021).

14 Liu, F. et al. Association Between Three Polymorphisms in BMAL1 Genes and Risk of Lung Cancer in a Northeast Chinese Population. DNA Cell Biol 38, 1437–1443, doi:10.1089/dna.2019.4853 (2019).

15 Qu, F. et al. Genetic polymorphisms in circadian negative feedback regulation genes predict overall survival and response to chemotherapy in gastric cancer patients. Sci Rep 6, 22424, doi:10.1038/srep22424 (2016).

16 Ye, Y. et al. The Genomic Landscape and Pharmacogenomic Interactions of Clock Genes in Cancer Chronotherapy. Cell Syst 6, 314–328 e312, doi:10.1016/j.cels.2018.01.013 (2018).

17 Liu, Z. et al. Dysregulation, functional implications, and prognostic ability of the circadian clock across cancers. Cancer Med 8, 1710–1720, doi:10.1002/cam4.2035 (2019).

18 Zhou, J. et al. The aberrant expression of rhythm genes affects the genome instability and regulates the cancer immunity in pan-cancer. Cancer Med 9, 1818–1829, doi:10.1002/cam4.2834 (2020).

19 Wu, Y., Tao, B., Zhang, T., Fan, Y. & Mao, R. Pan-Cancer Analysis Reveals Disrupted Circadian Clock Associates With T Cell Exhaustion. Front Immunol 10, 2451, doi:10.3389/fimmu.2019.02451 (2019).

20 Shilts, J., Chen, G. & Hughey, J. J. Evidence for widespread dysregulation of circadian clock progression in human cancer. PeerJ 6, e4327, doi:10.7717/peerj.4327 (2018).

21 Talamanca, L., Gobet, C. & Naef, F. Sex-dimorphic and age-dependent organization of 24-hour gene expression rhythms in humans. Science 379, 478–483, doi:10.1126/science.add0846 (2023).

22 He, H. et al. Combined analysis of single-cell and bulk RNA sequencing reveals the expression patterns of circadian rhythm disruption in the immune microenvironment of Alzheimer’s disease. Front Immunol 14, 1182307, doi:10.3389/fimmu.2023.1182307 (2023).

23 Liu, Y. et al. CRS: a circadian rhythm score model for predicting prognosis and treatment response in cancer patients. J Transl Med 21, 185, doi:10.1186/s12967-023-04013-w (2023).

24 Xu, Y. et al. Quantifying Personalized Shift-Work Molecular Portraits Underlying Alzheimer’s Disease through Computational Biology. J Prev Alzheimers Dis 11, 1721–1733, doi:10.14283/jpad.2024.161 (2024).

25 Anafi, R. C., Francey, L. J., Hogenesch, J. B. & Kim, J. CYCLOPS reveals human transcriptional rhythms in health and disease. Proc Natl Acad Sci U S A 114, 5312–5317, doi:10.1073/pnas.1619320114 (2017).

26 Duan, J. et al. tauFisher predicts circadian time from a single sample of bulk and single-cell pseudobulk transcriptomic data. Nat Commun 15, 3840, doi:10.1038/s41467-024-48041-6 (2024).

27 Hughey, J. J., Hastie, T. & Butte, A. J. ZeitZeiger: supervised learning for high-dimensional data from an oscillatory system. Nucleic Acids Res 44, e80, doi:10.1093/nar/gkw030 (2016).

28 Grabe, S., Ananthasubramaniam, B. & Herzel, H. Quantification of circadian rhythms in mammalian lung tissue snapshot data. Sci Rep 14, 16238, doi:10.1038/s41598-024-66694-7 (2024).

29. Hughes, M. E., Hogenesch, J. B. & Kornacker, K. JTK_CYCLE: an efficient nonparametric algorithm for detecting rhythmic components in genome-scale data sets. J Biol Rhythms 25, 372-380, doi:10.1177/0748730410379711 (2010).

30 Maas, M. B. et al. Circadian Gene Expression Rhythms During Critical Illness. Crit Care Med 48, e1294–e1299, doi:10.1097/CCM.0000000000004697 (2020).

31 Thaben, P. F. & Westermark, P. O. Detecting rhythms in time series with RAIN. J Biol Rhythms 29, 391–400, doi:10.1177/0748730414553029 (2014).

32 Deota, S. et al. Diurnal transcriptome landscape of a multi-tissue response to time-restricted feeding in mammals. Cell Metab 35, 150–165 e154, doi:10.1016/j.cmet.2022.12.006 (2023).

33 Wu, G., Anafi, R. C., Hughes, M. E., Kornacker, K. & Hogenesch, J. B. MetaCycle: an integrated R package to evaluate periodicity in large scale data. Bioinformatics 32, 3351–3353, doi:10.1093/bioinformatics/btw405 (2016).

34 Mei, W. et al. Genome-wide circadian rhythm detection methods: systematic evaluations and practical guidelines. Brief Bioinform 22, doi:10.1093/bib/bbaa135 (2021).

35 Consortium, G. T. et al. Genetic effects on gene expression across human tissues. Nature 550, 204–213, doi:10.1038/nature24277 (2017).

36 Verma, P. & Dalal, K. ADAMTS-4 and ADAMTS-5: key enzymes in osteoarthritis. J Cell Biochem 112, 3507–3514, doi:10.1002/jcb.23298 (2011).

37 Tortorella, M. D., Malfait, A. M., Deccico, C. & Arner, E. The role of ADAM-TS4 (aggrecanase-1) and ADAM-TS5 (aggrecanase-2) in a model of cartilage degradation. Osteoarthritis Cartilage 9, 539–552, doi:10.1053/joca.2001.0427 (2001).

38 Gossan, N. et al. The circadian clock in murine chondrocytes regulates genes controlling key aspects of cartilage homeostasis. Arthritis Rheum 65, 2334–2345, doi:10.1002/art.38035 (2013).

39 Wei, J. et al. Circadian rhythm disruption upregulating Per1 in mandibular condylar chondrocytes mediating temporomandibular joint osteoarthritis via GSK3beta/beta-CATENIN pathway. J Transl Med 22, 662, doi:10.1186/s12967-024-05475-2 (2024).

40 Szklarczyk, D. et al. The STRING database in 2023: protein-protein association networks and functional enrichment analyses for any sequenced genome of interest. Nucleic Acids Res 51, D638–D646, doi:10.1093/nar/gkac1000 (2023).

41 Aviram, R., Dandavate, V., Manella, G., Golik, M. & Asher, G. Ultradian rhythms of AKT phosphorylation and gene expression emerge in the absence of the circadian clock components Per1 and Per2. PLoS Biol 19, e3001492, doi:10.1371/journal.pbio.3001492 (2021).

42 Ruben, M. D. et al. A database of tissue-specific rhythmically expressed human genes has potential applications in circadian medicine. Sci Transl Med 10, doi:10.1126/scitranslmed.aat8806 (2018).

43 Altman, B. J. et al. MYC Disrupts the Circadian Clock and Metabolism in Cancer Cells. Cell Metab 22, 1009–1019, doi:10.1016/j.cmet.2015.09.003 (2015).

44 Mure, L. S. et al. Diurnal transcriptome atlas of a primate across major neural and peripheral tissues. Science 359, doi:10.1126/science.aao0318 (2018).

45 Jagannath, A., Taylor, L., Wakaf, Z., Vasudevan, S. R. & Foster, R. G. The genetics of circadian rhythms, sleep and health. Hum Mol Genet 26, R128–R138, doi:10.1093/hmg/ddx240 (2017).

46 Patke, A. et al. Mutation of the Human Circadian Clock Gene CRY1 in Familial Delayed Sleep Phase Disorder. Cell 169, 203–215 e213, doi:10.1016/j.cell.2017.03.027 (2017).

47 Devlin, P. F. Signs of the time: environmental input to the circadian clock. J Exp Bot 53, 1535–1550, doi:10.1093/jxb/erf024 (2002).

48 Roenneberg, T. & Merrow, M. The Circadian Clock and Human Health. Curr Biol 26, R432–443, doi:10.1016/j.cub.2016.04.011 (2016).

49 Colaprico, A. et al. TCGAbiolinks: an R/Bioconductor package for integrative analysis of TCGA data. Nucleic Acids Res 44, e71, doi:10.1093/nar/gkv1507 (2016).

50 Frankish, A. et al. Gencode 2021. Nucleic Acids Res 49, D916–D923, doi:10.1093/nar/gkaa1087 (2021).

51 Alboukadel, K., Marcin, K. & Przemyslaw, B. survminer: Drawing Survival Curves using ‘ggplot2*’*. (2021).

52. Terry, M. T. A Package for Survival Analysis in R. (2022).

53 Hadley, W. ggplot2: Elegant Graphics for Data Analysis. (Springer-Verlag New York, 2016).

54 Nan, X. ggsci: Scientific Journal and Sci-Fi Themed Color Palettes for ‘ggplot2’. (2023).

55 Thomas Lin, P. patchwork: The Composer of Plots. (2022).

56. Zuguang, G. Complex Heatmap Visualization. (2022).

57 Johan, L. {eulerr}: Area-Proportional Euler and Venn Diagrams with Ellipses. (2022).

58 Guangchuang, Y. ggplotify: Convert Plot to ‘grob’ or ‘ggplot’ Object. (2021).

